# Behavioral state-dependent lateralization of dorsal dentate gyrus c-Fos expression in mice

**DOI:** 10.1101/214239

**Authors:** Jake T. Jordan, M. Regis Shanley, Carolyn L. Pytte

**Author notes:** Correspondence should be addressed to: Jake Jordan, 65-30 Kissena Blvd, Psychology Department, Queens College, Flushing, NY 11367.

## Abstract

Hemispheric lateralization is a fundamental organizing principle of nervous systems across taxonomic groups with bilateral symmetry. The mammalian hippocampus is lateralized anatomically, physiologically, and chemically; however, functional asymmetries are not yet well understood. Imaging studies in humans have implicated the left and right hippocampus in specialized processing. However, it is not clear if lateralized activity occurs in the rodent hippocampus. c-Fos imaging in animals provides a measure of neuronal activity with a resolution at the level of single cells. The aim of this study was to determine whether lateralized activity-dependent c-Fos expression occurs in the rodent hippocampus. To understand functional lateralization of hippocampal processing, we compared interhemispheric expression of c-Fos in the dentate gyrus (DG), a structure involved in encoding new experiences, in mice that ran on a wheel, encoded a novel object, or remained in home cages. We found that wheel running induced the greatest amount of DG c-Fos expression in both hemispheres, with no difference between hemispheres. Object exploration resulted in left-lateralized DG c-Fos expression, whereas control mice were not lateralized. We then sought to determine whether differential consideration of hemispheres might influence the conclusions of a study by simulating common cell quantification methods. We found that different approaches led to different conclusions. These data demonstrate lateralization of neuronal activity in the mouse DG corresponding to the experience of the animal and show that differentially considering hemisphere leads to alternative conclusions.

## Introduction

Lateralization in the brain refers to interhemispheric asymmetry in the presence or dominance of a neural substrate [1]. Many forms of anatomical and physiological lateralization in the hippocampus have been well described; however, understanding functional hippocampal lateralization has been more elusive. In humans, both hippocampal hemispheres are recruited during spatial navigation, although activity in only the right hippocampus corresponds to navigation accuracy [2]. Conversely, the left entorhinal cortex appears to be specialized for object encoding [3] and the left hippocampus is specialized for object-cued spatial memory [4]. Spatial navigation involves actual or imagined self-motion through a real or virtual environment, whereas object memory consists of the processing of externally-originating sensory stimuli; invoking a long-held dichotomy in the characteristics of information integrated by the hippocampus described in the cognitive map theory [5].

Although direct functional comparisons cannot be made across paradigms, it seems that rodents likewise use both hemispheres for spatial memory yet also show hemispheric specializations in particular tasks. Spatial memory as tested on the Barnes Maze has been reported to be right-lateralized [6], while long-term spatial memory storage on the Y-Maze has been suggested to be left-lateralized [7, 8]. Interhemispheric asymmetries have also been reported in hippocampal physiology in mice [7, 9], suggesting lateralization in *how* information is processed, in addition to lateralization in which types of information are processed. Despite these data, one of the most crippling obstacles to translational hippocampal research is the widely held view that the rodent hippocampus does not exhibit the same inter-hemispheric differences that are seen in humans. Thus, a large majority of rodent hippocampal studies do not consider hemisphere.

Imaging immediate early gene expression (IEG) in animals provides several advantages over human imaging, such as single-cell resolution and the ability to examine activity during freely moving behaviors; however, no study has compared IEG expression in the left and right dorsal hippocampus (see [10], for ventral hippocampal imaging). Further, no study has examined lateralization in the dentate gyrus (DG), a structure important for the encoding of new hippocampus-dependent memories [11]. We examined how the behaviors of wheel running and exploration of a novel object are associated with IEG expression in the left and right dorsal DG. Interestingly, two studies that examined DG IEG expression but that differed in respect to treatment of hemispheres during cell quantification arrived at conflicting conclusions [12, 13]. Therefore, we simulated these and other common cell quantification approaches that are indifferent to hemisphere to determine whether such approaches may arrive at different conclusions.

## Methods and Experimental Design

This work was approved by the Queens College Institutional Animal Care and Use Committee (Protocol #177).

### Animals and Housing

Adult male (n=23) and female (n=22) C57Bl6 mice (Jackson Labs) were group housed (3-4) in standard shoebox plastic caging with bedding on a 12 hr light/dark cycle with free access to food and water.

### Behavior

Group-housed littermates were distributed roughly evenly across three treatment groups: 1) wheel running (WR, n= 5 males, 5 females), 2) exploration of a novel object in the home cage (OB, n= 6 males, 5 females), and 3) control home cage (CON, n= 9 males, 9 females). We sought to emphasize the relative salience of internally-generated motor information versus externally-originating environmental cues by training and testing the WR group in the dark to reduce visual input. In contrast, the OB mice were exposed to novel objects in the light. CON animals were housed in the dark for 24 hrs prior to sacrifice. In a follow-up experiment to test the effect of the difference in lighting between WR and OB groups, we determined c-Fos expression in a group that similarly experienced a novel object, but did so in the dark (OB-D, n=3 males, 3 females).

Behavioral training and testing were conducted in a small, dedicated behavior room. WR mice were trained to run on a wheel 24 hours prior to testing. For training, WR mice were individually brought to the behavior room in their home cage and within a few minutes placed in a cage identical to that of their home cage except containing a wheel. They were permitted 2 hours of housing in the wheel-cage, during which time they all ran to varying extents. They were then transferred to their home cage and dark housed in the behavior room overnight. The next day, WR mice were transferred from their home cage to the wheel-running cage, allowed 30 minutes of running, then transferred back to their home cage in the behavior room. Sixty-five minutes later, they were brought to a perfusion room and sacrificed. The entire procedure prior to transportation to the perfusion room (25 hr 35 min) was conducted in the dark.

OB mice were transported individually in their home cage to the behavior room and housed in the dark for 24 hours. The following day, the room light was turned on and a novel object (small PVC pipe) was placed in their home cage. After 30 minutes of exposure, the object was removed and the room light turned off. They remained in the dark for 65 minutes and then were transported to a perfusion room and sacrificed.

CON mice were transported individually in their home cages to the behavior room and housed undisturbed for 25 hr 35 min, time-matched to the behavioral experiments of the WR and OB groups. They were then transported to a perfusion room and sacrificed. Individual times spent wheel running and exploring the object were not quantified.

To determine whether the behavior room light contributed to differences in c-Fos expression between WR and OB groups, we repeated the object exploration treatment in an OB-D group in which the behavior room was dark throughout the experiment. Immunohistochemistry and c-Fos quantification of DG c-Fos expression for the OB-D mice was conducted separately from the first three groups and therefore was not included in the analysis of DG c-Fos+ cell densities across the other behavioral conditions.

### Immunohistochemistry

Seventy-five minutes following the end of behavior, mice were deeply anesthetized with 0.2 mL of Euthasol (Virbac) and perfused intracardially with 0.1 M phosphate buffered saline (PBS) and 4% paraformaldehyde in PBS. The brain was bisected and the dorsal hippocampus was coronally sectioned at 18 µm (two CON brains and one OB brain were cut at 30 µm) one hemisphere at a time. Cutting order of hemisphere was pseudorandomized. The brains of mice that explored an object in the dark were not bisected. Instead, one hemisphere was marked by removing a segment of ventral neocortex. Marked hemispheres were pseudorandomized. For all brains, every third section was mounted directly onto positively charged microscope slides with a random orientation in both axes such that hemisphere could not be determined by the section orientation. Slides were stored at −20°C until processed.

Tissue from WR, OB, and CON was processed together and distributed equally across immunohistochemistry batches. The OB-D group was processed separately. Frozen sections were rinsed once in 0.1 M PBS for 10 minutes. A block containing 10% donkey serum and 3% Triton-X in phosphate buffer (PB) was applied for 30 minutes, followed by a 48-hour incubation in anti-c-Fos made in rabbit (Millipore, ABE457, 1:1000) in block. Sections were then rinsed in 0.1 M PBS three times for 10 min each and then incubated with Cy3-conjugated donkey anti-rabbit for 2 hours (Millipore, AP182C, 1:200) in block or PB/S). Sections were rinsed in 0.1 M PBS three times for 10 min each before a serial dehydration in ethanols followed by immersion in xylene and cover slipping with Krystalon (Harleco).

### Cell Quantification

Microscopy was conducted blind to hemisphere for all mice. Microscopy was conducted blind to treatment for groups WR, OB, CON, but aware of treatment for OB-D as this group was processed independently after the others. The DG was traced under 4x magnification using Neurolucida software (Microbrightfield) coupled to an Olympus BX51 microscope. Cy3-labeled cells within the DG were counted under 60x magnification using a rhodamine filter. For brains sectioned at 18 µm, 3-5 sections (54-90 µm) were quantified per hemisphere per mouse; for brains sectioned at 30 µm, 2-4 sections (60-120 µm) were quantified per hemisphere per mouse. Amount of tissue sampled did not correlate with c-Fos+ cell counts. Densities of c-Fos expression within hemispheres were calculated by determining the number of c-Fos+ cells within the DG boundary divided by the volume sampled. We analyzed our tissue comparing left and right c-Fos expression across groups to determine whether c-Fos expression was lateralized, and if so, whether it changed with experience.

Because the amount of c-Fos expression may vary more widely among individuals than between hemispheres, we also calculated a lateralization index (LI), which shows hemispheric asymmetry normalized by the total number of c-Fos+ cells in each animal and compared LI values across groups. Positive values indicate more c-Fos+ cells in the left hemisphere relative to the right and negative values indicate more cFos+ cells in the right hemisphere.

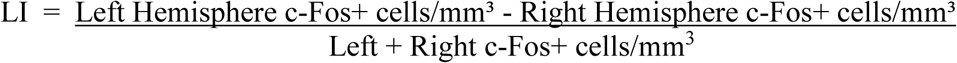

In the course of this work, we found that hemisphere use varies greatly in the literature and we became interested in the potential effect of hemisphere selection on the outcome of hippocampal studies more broadly. Therefore, we compared c-Fos expression across treatment groups using the left hemisphere only, the right hemisphere only, and both hemispheres combined to determine whether hemisphere use affected the results.

We further simulated commonly used cell quantification methods that ignore possible hemispheric asymmetries, by conducting three additional computations. To model experiments that look at expression unilaterally, we compared cell counts across treatments using primarily one hemisphere and occasionally the other, mimicking occasions of tissue loss. We then modeled experiments that quantify cells unilaterally, but do not prefer one hemisphere or another. To do this, we performed 1,000 simulations for each of the three conditions in which: 1) 80% of quantifications were taken from the left hemisphere and 20% were taken from the right; 2) 20% of quantifications were taken from the left hemisphere and 80% were taken from the right and 3) 50% of quantifications were taken from the left hemisphere and 50% were taken from the right hemisphere. Within these restrictions, all selections of hemisphere were random across mice. Between simulations, all draws were random with replacement.

### Statistics

#### Sex differences

A two-way ANOVA was used to compare bilateral DG c-Fos expression between sexes and across behavioral groups. A post-hoc power analysis was performed on each group to determine the sample size needed to reach a power level of 0.8 at a confidence level of 0.05.

#### Lateralization of c-Fos expression within treatment groups

We compared the numbers of cells expressing c-Fos between the left and right DG within each treatment group using paired t-tests.

#### Lateralization of c-Fos expression across WR, OB, and CON groups

A two-way mixed-model ANOVA, with Tukey’s post-hoc test, was used to compare c-Fos in the left and right hemisphere (repeated measure) between treatments. The OB-D group could not be included as the tissue was processed independently from that of the other treatments.

#### Lateralization indices of c-Fos expression across WR, OB, CON, and OB-D groups

A one-way ANOVA was used to determine whether lateralization indices differed across treatments. This was followed by a Dunnett’s multiple comparisons post-hoc test comparing the lateralization index of each behavioral treatment to that of CON mice only.

#### Comparative cell quantification methods

We used one-way ANOVAs and Tukey’s post-hoc tests to compare c-Fos expression across treatment groups using data from a single hemisphere, both hemispheres combined, and random combinations of hemispheres in 80:20, 20:80 or 50:50 left:right ratios.

## Results

### There were no sex differences in c-Fos expression

To determine whether c-Fos expression differed in males and females, we compared bilateral DG c-Fos expression across behavioral groups and between sexes. Behavioral condition had a significant effect on expression (F(2,33)=26.12, P<0.0001). There was no effect of sex on expression (F(1,33)=0.616, P=0.438) and there was no interaction between behavior and sex (F(2,33)=0.289, P=0.751). However, it is possible that sex differences were not detected due to low sample sizes. We performed a power analysis on each behavioral group to detect sample sizes needed to reach a power level of 0.8 at a confidence level of 0.05 and found this would require a total of 85 mice in the CON group, 9,812 mice in the WR group and 46 mice in the OB group. Thus, we combined males and females for all subsequent analyses.

### C-Fos expression was lateralized in OB mice

Our first goal was to determine whether activity, measured by c-Fos expression, was lateralized between hemispheres within each of the behavioral contexts. We found that within CON or WR mice, numbers of c-Fos expressing cells did not differ between hemisphere (CON: df 17, t=-1.539, P=0.142; WR: df 9, t=0.086, P=0.934, paired t-test). However, there were significantly more c-Fos expressing cells in the left than in the right hemisphere within the OB group (df 10, t=2.681, P=0.023, paired t-test).

### Object exploration, but not wheel running, lateralized DG c-Fos expression to the left hemisphere

We were also interested in whether there were hemispheric differences in c-Fos expression after engaging in wheel running or exposure to a novel object compared to the context of standard-housing. We found a main effect of behavioral condition (F(2,36)=27.39; P<0.0001) but not of hemisphere (F(1,36)=0.282; P=0.599). There was an interaction between these two factors (F(2,36)=3.357; P=0.040, two-way mixed ANOVA, Figure 1A). Post-hoc analyses revealed that c-Fos expression in the left hemisphere of OB and WR mice differed from that of CON mice (OB vs. CON: P=0.014; WR vs. CON: P<0.0001, Tukey’s test) and left hemispheres in OB and WR mice differed from each other (OB vs. WR: P=0.0003, Figure 1A, Tukey’s test). Across right hemispheres, WR mice had greater expression than either CON or OB mice (WR vs. CON: P<0.0001; WR vs. OB: P<0.0001, Tukey’s test). However, unlike in the left hemisphere, right hemisphere expression of c-Fos did not differ between OB and CON mice (P=0.418, Figure 1A, Tukey’s test).

**Figure 1.**
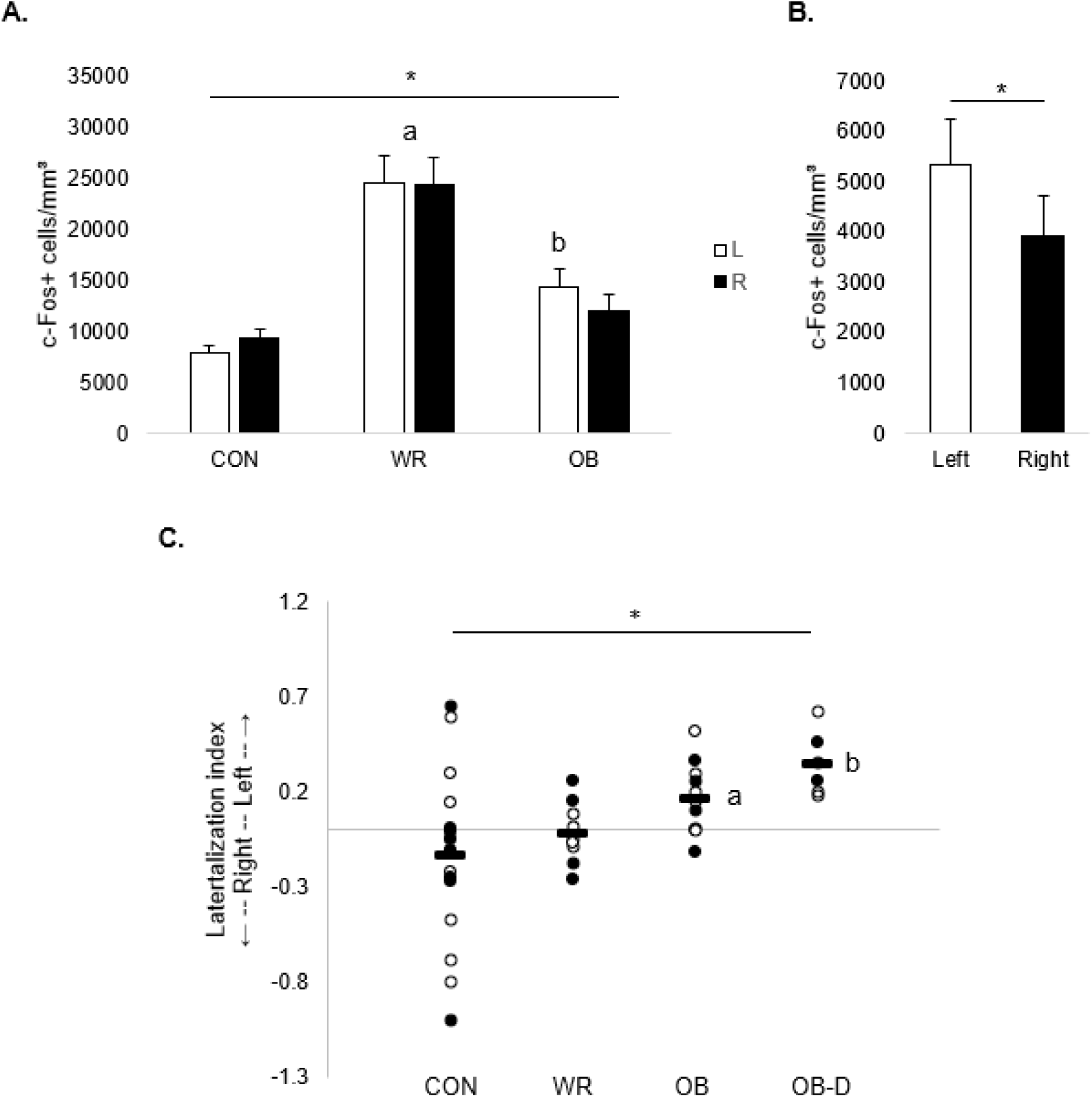
Exposure to a novel object preferentially recruited the left DG, while wheel running equally recruited both DG. (A) Dorsal DG c-Fos expression across hemispheres and behavioral groups showed an overall effect of wheel running on left- and right-hemisphere c-Fos expression and an interaction between treatment group and hemisphere. * indicates main effect, P<0.05; *a:* post-hoc tests indicate that c-Fos+ cell counts in both the left and right hemispheres of the WR group were greater than those in either the CON or OB groups; *b:* indicates an effect of object exploration on left-hemisphere c-Fos expression compared to the left hemisphere of controls; letters indicate P<0.05. (B) Left and right densities of DG c-Fos expression in OB-D mice. * indicates a difference between hemispheres, P<0.05. (C) Degree of hemispheric lateralization for individuals in each behavioral group. * indicates a main effect of treatment group, P<0.05; *a, b:* post-hoc tests show that lateralization indices in OB and OB-D differ from that of CON. Males are shown as open circles and females are shown as closed circles; group means are shown as horizontal black bars.

To determine whether room lighting contributed to the lateralization of DG c-Fos expression in OB mice, we examined DG c-Fos expression in OB-D mice, which underwent the same behavioral treatment as OB mice, but in the dark (as were CON and WR mice). Like the OB group, OB-D mice had significantly higher DG c-Fos expression in the left as compared to the right hemisphere (df 5, t=6.401, P=0.001, paired t-test, Figure 1B). Overall levels were lower in OB-D than in the other groups; however, we cannot determine whether this was due to the treatment or differences in tissue processing as these mice were processed separately from the other groups.

Finally, we used a lateralization index to compare relative c-Fos expression across hemispheres normalized to individual expression levels. We found a significant difference in the lateralization index across treatment groups (df 3, F=4.611, P=0.007, one-way ANOVA, Figure 1C). Post-hoc comparisons indicated that the lateralization indices of OB-D and OB mice differed significantly from that of CON mice and there was no difference between WR and the CON group (CON vs. OB: P=0.006; CON vs. OB-D: P=0.006; CON vs. WR: P=0.678, Dunnett’s post-hoc test).

### Widely-used cell quantification methods produced different results

A majority of hippocampal cell quantification studies do not account for potential hemispheric asymmetries and either pool hemispheres, reporting total cells across hemispheres, or only use one hemisphere. In the latter case, the hemisphere used may be consistent (i.e. the left or the right hemisphere is quantified for each animal) or it may not be identified (cell counts may include the left hemisphere of some subjects and the right hemisphere from others, or cell counts may include mixed hemispheres from some or all subjects).

To address potential differences in outcomes corresponding to hemisphere selection, we first performed one-way ANOVAs on c-Fos expression in each hemisphere individually. When including only the left hemisphere in our analysis, we found an effect of behavioral condition on DG c-Fos expression (F(2,36)=25.37, P<0.0001, one-way ANOVA). Post-hoc tests revealed significant differences among all three behavioral groups (OB vs. CON: Q=3.96, P=0.022; WR vs. CON: Q=10.07, P<0.01; OB vs. WR: Q=5.62, P<0.01, Tukey’s post-hoc test, Figure 2A). When including only the right hemisphere in our analysis, we again found an effect of behavioral condition on DG c-Fos expression (F(2,36)=24.91, P<0.0001, one-way ANOVA). In this analysis, WR mice had higher c-Fos expression than both CON (Q=9.80, P<0.01) and OB mice (Q=7.22, P<0.01), however, there was no difference between OB and CON mice (Q=1.85, P=0.400, Tukey’s, Figure 2B).

**Figure 2.**
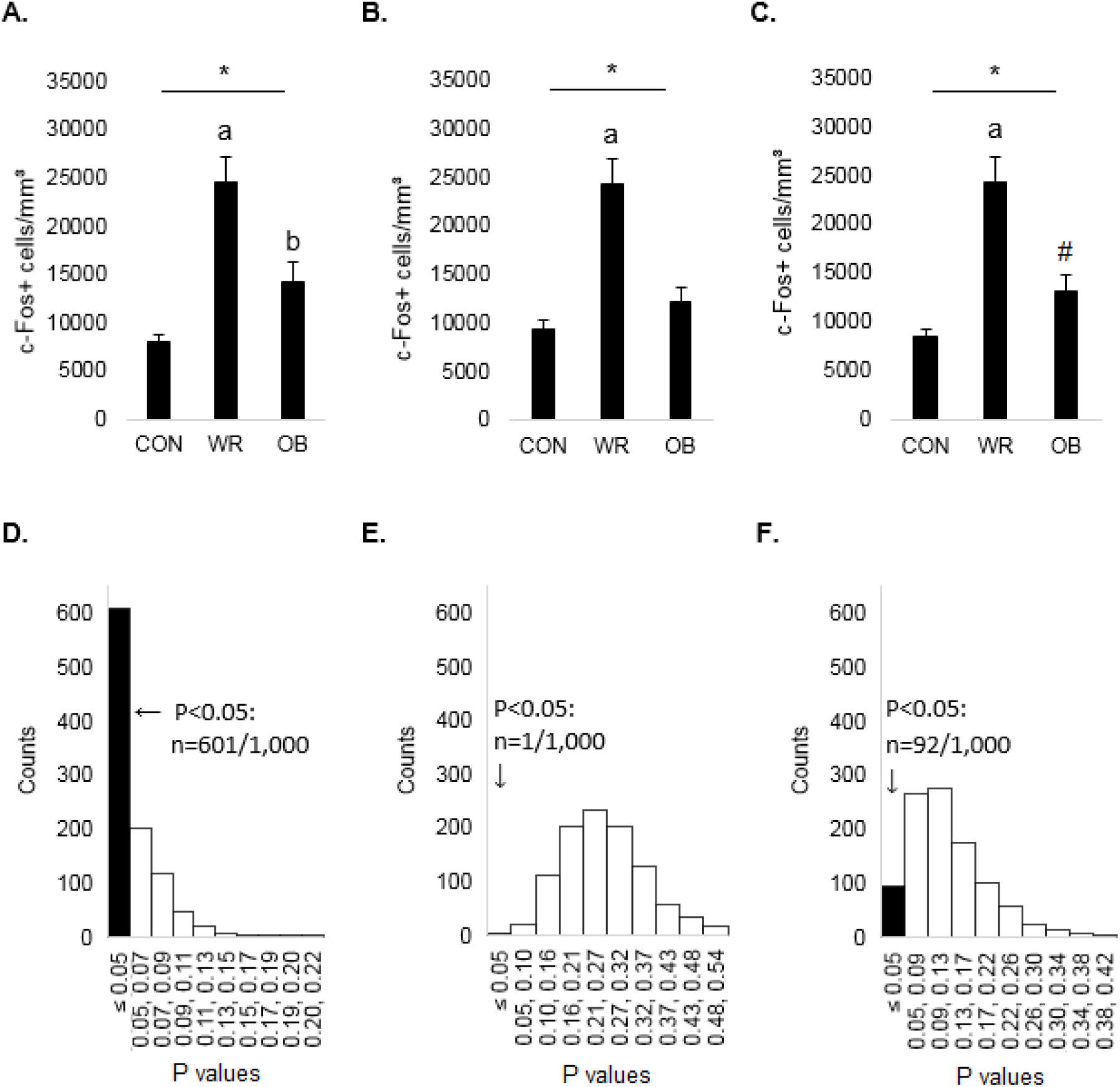
Ignoring hemispheres during cell quantification can lead to different conclusions. (A) DG c-Fos expression in the left hemisphere only. (B) DG c-Fos expression in the right hemisphere only. (C) DG c-Fos expression when hemispheres were pooled bilaterally. (D) Instances of P values determined by Tukey’s post-hoc tests comparing OB and CON, conducted after each of 1,000 iterations of random combinations using 80% of our c-Fos data from the left hemisphere and 20% from the right hemisphere. Binned ranges of P values are shown along the x axis and instances of P values per bin are shown along the y axis. (E) Instances of P values determined by Tukey’s post-hoc tests comparing OB and CON, conducted after each of 1,000 iterations of random combinations using 20% of our left and 80% of our right hemisphere c-Fos data. (F) Instances of P values determined by Tukey’s post-hoc tests comparing OB and CON, conducted after each of 1,000 iterations of random combinations using 50% of our left and 50% of our right hemisphere c-Fos data. Black bars in D, E, F indicate the number of instances in which P values were less than 0.05. * indicates a main effect of behavioral group, P<0.05; *a:* indicates a significant difference between WR vs. OB and WR vs. CON, P<0.05; *b:* indicates a significant difference between OB vs. CON, P<0.05; # indicates a trend toward a significant difference between OB vs. CON, P=0.088.

We then analyzed DG c-Fos expression bilaterally. For each mouse, we summed cell counts across hemispheres and divided by the summed volume sampled in both hemispheres, producing a measure of bilateral c-Fos density. This analysis yielded an effect of behavioral condition on bilateral DG c-Fos expression (F(2,36)=27.49, P<0.0001; one-way ANOVA). Post-hoc analysis revealed significantly greater expression in WR mice compared to CON (Q=10.45; P<0.01; Tukey’s; Figure 2C) and OB mice (Q=6.73, P<0.01; Tukey’s; Figure 2C). There was a trend toward a significant difference in expression in OB and CON mice (Q=3.09, P=0.088; Tukey’s; Figure 2C).

Next, we simulated experiments in which the same hemisphere was not consistently quantified for each animal in a group. The results of these simulations did not differ in the outcome of the ANOVAs; all showed a significant main effect of treatment (P <0.05) and no effect of hemisphere. They also all resulted in significant pairwise comparisons between the WR group and both CON and OB (Tukey’s post-hoc tests). However, the outcomes did vary in whether the post-hoc tests determined that the means of OB and CON groups differed. In our first simulation, we used values of c-Fos expression in the left hemisphere in 80% of the animals, and c-Fos expression in the right hemisphere in the remaining 20% of the animals within each treatment. In 601 out of 1,000 simulations, a post-hoc test between OB and CON mice produced P < 0.05 (Figure 2D). We then simulated experiments with the reverse ratio in which c-Fos expression in the left hemisphere was used in 20% of the animals, and expression in the right hemisphere was used in the remaining 80% of animals. Strikingly, in only one out of 1,000 of these simulations was a significant difference detected between OB and CON mice (Figure 2E). Finally, we simulated experiments in which we used 50% of our c-Fos values from the left and 50% of our c-Fos values from the right hemispheres. In 92 of 1,000 simulations there was a significant difference between OB and CON mice (Figure 2F).

## Discussion

We quantified neuronal activity-related c-Fos expression in the left and right DG following wheel running, object exploration, and home cage housing. Wheel running in the dark upregulated c-Fos expression bilaterally and showed no hemispheric asymmetry. Interestingly, exploration of a novel object with the room lights on upregulated c-Fos expression in the left, but not the right hemisphere compared with controls housed in the dark. With the room lights off, mice similarly exposed to an object showed lower DG c-Fos expression than the controls, however, the tissue for the OB-D mice was processed and expression quantified separately (and using a primary antibody from a different lot), and thus, we cannot directly compare expression levels of this group across treatments. More importantly for our objectives, object exploration in either the light or dark resulted in left-dominant DG c-Fos expression compared to controls.

Although the rodent hippocampus has generally been considered to be functionally symmetric, accumulating evidence suggests that there are indeed rodent hippocampal functional asymmetries (see [14] for review). The assumed lack of hemispheric specialization in rodents has in part been based on substantial interhemispheric connectivity between the two hippocampi as subfields send both ipsi- and contralateral projections. In contrast, it is widely believed that humans have only ipsilateral projections from CA3 to CA1 [15]. Although the presence of bilateral projections in humans has been argued for [16, 17, 18], this position has not been widely adopted [19]. Rather, it has been suggested that in humans and other primates, hemispheric isolation due to the absence of interhippocampal pathways has resulted in a higher degree of hemispheric processing specialization than in other mammals [20]. However, the anatomy of inter-hemispheric projections need not require or predict functional asymmetry.

A recent study showed that sharp-wave ripples in the rat hippocampus, thought to function in memory consolidation, have coordinated activity within either, but not across, hemispheres [21]. This suggests that even though information may be shared across hemispheres in rodents, cell assemblies corresponding to a particular experience may be intrahemispheric and their information consolidated in isolation to each other. This is consistent with other work indicating that spatial memory may be lateralized to one hemisphere or the other ([22], rats; [6, 7. 8], mice). Further, neurophysiological studies in mice have identified lateralization in some of the key neural substrates, such as LTP and NMDA receptor subunits, thought to underlie hippocampal function [7, 9, 23, 24, 25]. Taken together, this suggests that despite potential differences in direct interhemispheric communications between humans and rodents, hippocampal hemispheric specialization may be a shared property.

It is not yet clear whether rodent and human hippocampal lateralization show the same patterns of functional division of labor. In humans, a role of the right hippocampus in allocentric spatial navigation has been widely documented [2, 26, 27, 28, 29] whereas a clear understanding of left hippocampus processing remains more elusive. The role of the left side has been suggested to relate to the transfer of recently formed memories to long term storage (in rats, [22]) or function in autobiographical memory formation (in humans, [30]). More recently, the left hippocampal formation in humans has been shown to be preferentially active during an object recognition task [3, 4] whereas the right hippocampus showed greater response in a place task [4]. It is intriguing that we found similar left side preference for object processing in mice, suggesting perhaps greater functional parallels between humans and rodents than has been recognized. However, it is important to note that we do not know whether the lateralization evoked by the OB exposure is specific to this experience with the object or whether it reflects a more general processing function of the hippocampus. Additional behavioral tasks would help clarify this issue.

In addition, we found that the use of common cell quantification methods resulted in different conclusions regarding the effects of experience on neuronal activity in the rodent hippocampus. In this context, it is intriguing that two similar studies examining IEG expression in the rodent DG arrived at contradictory conclusions. Deng et al. [12] found that re-exposure to a fear-conditioned context did not reactivate DG neurons that were active during the conditioning trial. However, Denny et al., [13] found that re-exposure to a fear-conditioned context in a similar task paradigm did in fact reactivate the same population of neurons that were active during initial encoding. The two studies used different genetic mouse models, which may have contributed to their differing results. However, the two reports also differed in their approaches to cell quantification with respect to hemisphere. Deng et al. [12] quantified unilaterally, primarily using the right hemisphere for all mice, but on occasion using the left hemisphere if the right was damaged (personal communication). When primarily using the right hemisphere, our data set led to the finding that DG c-Fos expression was not different between OB and CON mice, while primarily using the left hemisphere lead to the finding that DG c-Fos expression was indeed significantly different between the same groups. Denny et al. [13] pooled hemispheres and quantified bilaterally (personal communication). This approach in our study led to a trend (P=0.088) when comparing OB and CON DG c-Fos expression. Thus, considering hemispheric differences in rodent models may potentially resolve discrepancies in reported findings. Moreover, considering hemisphere may in addition provide critical insight into the mechanisms of hippocampal function.

## Acknowledgements

We thank Jeff Beeler and Daniel McCloskey for input on experimental design.

## Author Contributions

JTJ designed the study. JTJ performed the experiments. JTJ, MRS, and CLP analyzed the data and wrote the manuscript.

## Data Accessibility

All data for this manuscript will be made available upon email request.

## Abbreviations

CON: control
DG: dentate gyrus
IEG: immediate early gene (IEG)
LTP: long-term potentiation
NMDA: N-methyl-D-aspartate
OB: Object-exploration
OB-D: Object exploration in the dark
PB: phosphate buffer
PBS: phosphate buffered saline
WR: Wheel running

